# Single-cell glycolytic activity regulates membrane tension and HIV-1 fusion

**DOI:** 10.1101/693341

**Authors:** Charles A. Coomer, Irene Carlon-Andres, Maro Iliopoulou, Michael L. Dustin, Ewoud B. Compeer, Alex A. Compton, Sergi Padilla-Parra

## Abstract

There has been resurgence in determining the role of host metabolism in viral infection yet deciphering how the metabolic state of single cells affects viral entry and fusion remains unknown. Here, we have developed a novel assay multiplexing genetically encoded biosensors with single virus tracking (SVT) to evaluate the influence of global metabolic processes on the success rate of virus entry in single cells. We found that cells with a lower ATP:ADP ratio prior to virus addition were less permissive to virus fusion and infection. These results indicated a relationship between host metabolic state and the likelihood for virus-cell fusion to occur. SVT revealed that HIV-1 viruses were arrested at hemifusion in glycolytically-inactive cells. Interestingly, cells acutely treated with glycolysis inhibitor 2-deoxyglucose (2-DG) become resistant to virus infection and also display less surface membrane cholesterol. Addition of cholesterol in these in glycolytically-inactive cells rescued the virus entry block at hemifusion and enabled completion of HIV-1 fusion. Further investigation with FRET-based membrane tension and membrane-order reporters revealed a link between host cell glycolytic activity and host membrane order and tension. Indeed, cells treated with 2-DG possessed lower plasma membrane lipid order and higher tension values, respectively. Our novel imaging approach that combines lifetime imaging (FLIM) and SVT revealed not only changes in plasma membrane tension at the point of viral fusion, but also that HIV is less likely to enter cells at areas of higher membrane tension. We therefore have identified a connection between host cell glycolytic activity and membrane tension that influences HIV-1 fusion in real-time at the single-virus fusion level in live cells. As glycolytic activity sets membrane tension levels by altering cellular cholesterol surface levels, our results suggest additional previously unknown benefits of cholesterol-lowering medication in HIV-1 infection.

## INTRODUCTION

It is well-established that HIV-1 preferably infects target cells by engaging its gp160 trimer of heterodimers (i.e. gp120 and gp41) with host receptor CD4 (Dalgliesh et al. 1984) and co-receptors CCR5 (Dragic et al. 1996) or CXCR4 (Feng et al. 1996). Subsequently, the receptor and co-receptor interactions trigger conformational changes in the gp41 transmembrane subunit, facilitating virus-cell fusion and the formation of a post-fusion, thermodynamically-stable six-helix bundle (Gallo et al. 2003). Identifying elements that underpin the earliest stages of HIV-1 infection is a priority in the search for an HIV cure (Deeks et al. 2016). Furthermore, numerous studies have highlighted the importance of cellular metabolism in T cell-mediated antiviral responses and control of viral infection (Buck et al. 2017; Pallett et al. 2019). However, the role played by single-cell metabolic activity in susceptibility to HIV-1 invasion has not been addressed.

Upon activation, T cells increase rates of oxidative phosphorylation and glycolysis to cope with higher energy demands associated with immune functions (Pearce et al. 2013). Expression of the glucose transporter Glut1 is increased upon T cell activation and is required for post-entry HIV replication (Loisel-Meyer et al. 2012). Moreover, the Glut1-associated increased glycolytic flux has also been observed in HIV-1 infected cells (Hollenbaugh et al. 2011, 2016) and affects virus pathogenicity and production (Hegedus et al. 2014). More recently, it has been shown that basal glycolytic activity of CD4^+^ T cell subsets correlates with susceptibility to HIV-1 infection (Valle-Casuso et al. 2018). The authors showed that chronic, suboptimal inhibition of glycolytic activity impairs HIV-1 infection in T cells, but it was unclear which stage of HIV-1 infection was affected (Valle-Casuso et al. 2018). Pyruvate made from glycolysis may enter the citric acid cycle to yield biosynthetic intermediates (such as oxaloacetate and citrate) and reduced electron carriers (NADH and FADH_2_). Although in activated T cells most pyruvate is excreted as lactate, still increased cytosolic levels of citrate and NAD^+^ are found (Buck et al. 2015) suggesting that this pathway is still in use by activated T cells. Citrate is a precursor of cytosolic acetyl-CoA which is required for cholesterol biosynthesis and a major determinant of cellular membrane architecture (Liu et al. 2013).

Cholesterol is required for the formation and maintenance of liquid ordered (Lo) membrane domains in addition to reducing membrane tension (Feigenson 2006; Biswas et al. 2019). Higher plasma membrane cholesterol content stretches lipid tails in Lo domains, a discontinuity within the bilayer is created at the phase boundary between Lo and Ld domains, manifesting in line tension that is directly implicated in regulating membrane curvature. Hemifusion is regulated by membrane curvature—positive curvature negates the propensity of fusion pores to progress to full fusion, whereas negative curvature facilitates this process (McMahon and Gallop 2005; Chernomordik and Kozlov 2008). Consequently, increased cholesterol establishes line tension between Lo and liquid disordered (Ld) domains and generates negative spontaneous curvature in support of fusion, whereas cholesterol depletion increases membrane tension, decreases line tension between Lo and Ld domains and enables positive spontaneous membrane curvature to disfavor fusion (Baumgart et al. 2003; Yang et al. 2016; Morris and Homann 2001; Biswas et al. 2019; Tsai and Feigenson 2019).

Membrane order, tension and curvature are significant contributors to several membrane-regulated processes in human cells important for viral infection, particularly virus entry (i.e. fusion and endocytosis) and egress (Ewers and Helenius 2011; Kozlov and Chernomordik 2015). In addition, boundaries between cholesterol-enriched Lo and Ld microdomains are well-established determinants of successful entry for a range of viruses, including HIV-1 (Campbell et al. 2001; Yang et al. 2015, 2016, 2017). The phase boundaries between Lo and Ld domains are thought to represent the site of HIV-1 entry, which may be due to line tension derived from Lo and Ld boundaries which drive membrane bending to facilitate fusion (Yang et al. 2015, 2016). Intriguingly, it has been demonstrated that plasma membrane cholesterol content of dendritic cells, macrophages, and T cells in HIV-1-infected long-term non-progressors is substantially reduced compared to that of rapid progressors, and that the cellular cholesterol content correlates with the capacity of HIV-1 to spread (Rappocciolo et al. 2014; Delucia and Rinaldo 2018). Therefore, an unexplored relationship may exist between the host cell metabolic state, cholesterol, and virus entry into cells.

A significant development in the study of cellular metabolism is the application of fluorescence-lifetime imaging microscopy (FLIM) in individual cells (Becker 2012; Suhling et al. 2015). FLIM provides resolution in single cell microenvironments that are missed by population-based or spectral methods. Here, we observed higher rates of HIV-1 fusion and infection in cells with high glycolytic activity, as reported by their ATP:ADP ratios and intracellular lactate concentrations, compared to cells with low glycolytic activity. Further analysis demonstrated that targeted inhibition of glycolysis by 2-deoxy-d-glucose (2-DG) drastically decreased HIV-1 fusion and infection. Multicolour real-time single virus tracking (SVT) revealed that HIV-1 entry is arrested at the stage of hemifusion in cells that are glycolytically inactive. 2-DG-treated cells displayed a loss of membrane cholesterol and addition of water-soluble cholesterol to these cells rescued HIV-1 fusion. Moreover, 2-DG treatment decreased membrane order and increased tension of cells as measured by FRET-based biosensor probes using FLIM. To further investigate the connection between membrane tension and HIV-1 entry, we developed a novel assay that combines FLIM and single virus tracking techniques to visualize and identify factors pivotal to HIV-1 infection at high-spatiotemporal resolution in single cells. Through this approach, we initially show that HIV-1 particles generate increased flux in global host-cell membrane tension that is, at least in part, CCR5-dependent. Furthermore, we illustrate that HIV-1 requires local decreases in membrane tension during the process of virus-cell fusion, which is prevented during the inhibition of glycolysis, yet rescued via cholesterol supplementation. Overall, our results demonstrate a functional link between the glycolytic activity of the host cell and the success of HIV-1-mediated virus-cell fusion, with an important role for membrane cholesterol, establishing a favourable physical environment which enables mechanical alterations in the host membrane to occur to enable virus entry.

## RESULTS

### Basal Metabolic State is a Determinant of HIV-1 Infection in T cells and Reporter Cells

There is an inherent hierarchy in the susceptibility of CD4 T cell subsets to HIV-1 infection. Recent work observed that overall T cell subset metabolic activity correlates with susceptibility to replication by HIV pseudotyped with VSV-G envelope protein (HIV-1_VSV-G_-eGFP) (Valle-Casuso et al. 2018). Here, we use TZM-bl and MT4 T cells that are more homogeneous in metabolic state and have high glycolytic activity similar to activated CD4 T cells susceptible to HIV-1 infection (Thompson 2017). Both TZM-bl and MT4 T cells are highly susceptible to HIV-1_NL4.3_ and HIV-1_VSV-G_ infection (Supplemental Figure 1A). Importantly, TZM-bl cells report HIV-1 infection with high sensitivity by β-Gal. Initially, we observed a similar dependence of HIV-1 infection on glycolytic activity in MT4 T cells treated acutely (i.e. 2 hours) with 2-DG prior to infection with either HIV-1_VSV-G_ (Figure 1A). Furthermore, we determined that acute arrest of aerobic respiration with oligomycin, an inhibitor of the F_0_ proton channel of ATP synthase, led to no such decrease in HIV-1_VSV-G_ infection (Figure 1B). Importantly, when treating T cells with HIV-1_NL4.3,_ we observed a similar dependence between 2-DG treatment and inhibition of virus infection (Supplemental Figure 1A).

**Figure 1.**
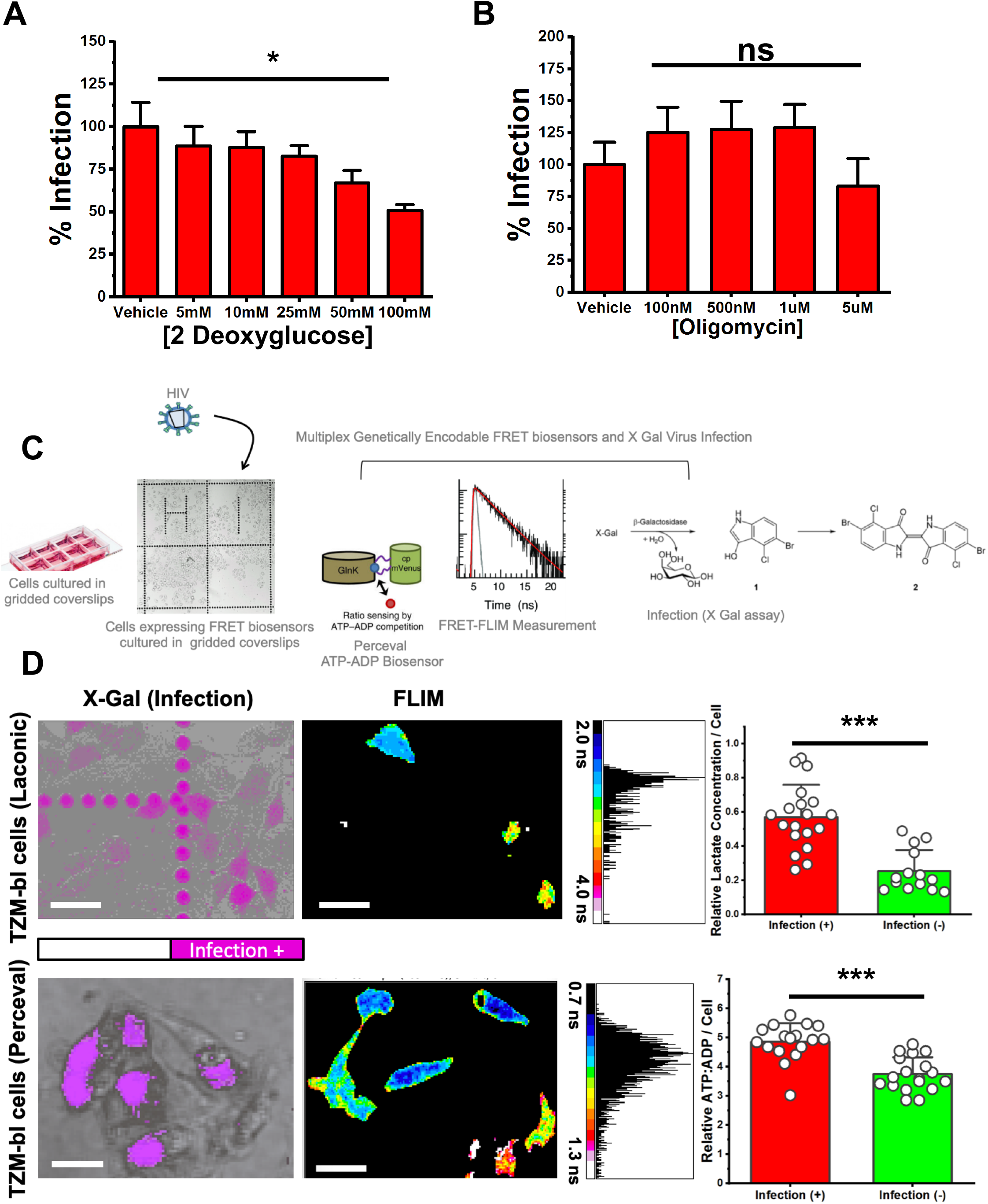
Relative lactate and ATP/ADP concentrations in single cells correlate with HIV-1 infection. A.) Bar charts depicting % eGFP-expressing cells as a marker of infection illustrating that acute treatment with 2-DG led to reductions in HIV-1_VSV-G_ infection in human MT4 T cells. B.) Bar charts depicting % eGFP-expressing cells as a marker of infection illustrating that acute treatment with increasing concentrations of oligomycin did not inhibit HIV-1_VSV-G_ infection in human MT4 T cells. C.) Approach to evaluate if basal, single-cell ATP:ADP ratio could predict HIV-1_JR-FL_ infection in TZM-bl cells. TZM-bl cells were seeded onto an Ibidi® 8-well gridded dish and transiently transfected with metabolic sensing biosensors Laconic or Perceval. FLIM images of cells transiently expressing these biosensors were recorded, denoting the location of the cells on the grid. These cells were then treated with HIV-1_JR-FL,_ and after 48 hours a β-galactosidase assay was recorded on the same cells as a readout of infection. D.) Representative images of β-galactosidase recorded 48 hours after HIV-1_JR-FL_ infection (left) and basal FLIM images of intracellular lactate biosensor Laconic-expressing TZM-bl cells taken before infection (right) illustrating cells with higher intracellular lactate concentrations (warm colours) were more likely to be infected by HIV-1_JR-FL_ (top row). Representative images of β-galactosidase recorded 48 hours after HIV-1_JR-FL_ infection (left) and basal FLIM images of intracellular ATP:ADP biosensor Perceval-expressing TZM-bl cells taken before infection (right) illustrating cells with higher intracellular ATP:ADP concentrations (cool colours) were more likely to be infected by HIV-1_JR-FL_ (bottom row). *** p<0.001 as determined by Student T test of three independent experiments.

To determine how a cell’s metabolic state correlates to HIV-1 infection, we employed fluorescence lifetime imaging microscopy (FLIM) which enables the quantification of the activity of metabolite reporters at the single cell level. We use FLIM as this methodology provides measurements that are independent of a fluorophore’s concentration, independent of pH (Supplemental Fig 1F) (Tantama et al. 2011), and increases the reporter’s dynamic range (Supplemental Figure 1B-E) (Szmacinski and Lakowicz 1995). Our general strategy to determine if metabolic states could influence virus entry and subsequent infection is depicted in Figure 1C. Briefly, we transiently transfected Perceval (Berg et al. 2009), an established ratiometric reporter of the ATP:ADP ratio, or Laconic (San Martín et al. 2013), an intracellular lactate biosensor to monitor glycolytic flux in TZM-bl cells via FLIM. This approach allowed us to determine if relative ATP:ADP ratios or relative lactate concentrations prior to infection could be predictive of cellular permissiveness to HIV-1 infection, as measured by β-galactosidase activity (Smale 2010). The seeding of cells onto gridded coverslips enabled single cells to be tracked over time. Interestingly, cells that possessed higher relative lactate concentrations or a higher relative ATP:ADP ratio were more likely to become infected by HIV-1_JR-FL_ (Figure 1D, top row) (Figure 1D, bottom row). This shows that the correlation between glycolytic activity and HIV-1 infection holds true not only in drug-induced differences but also in the small physiological differences within live cells.

### Basal Metabolic State is a Determinant of HIV-1 Fusion in Reporter Cells

The separation between HIV-1_JR-FL_-infected and non-infected cells based on their ATP:ADP ratios and relative lactate concentrations in reporter cells prior to infection prompted us to determine the stage at which HIV-1 infection is inhibited. To determine if the preference for HIV-1 to infect glycolytically-active cells is based on pre- or post-fusion events in cells, we multiplexed our FLIM-based reporters with the β-lactamase assay (BlaM) (Jones and Padilla-Parra 2015) (Figure 2A), which provides a readout of viral access to cytoplasm (virus-cell fusion). Importantly, the BlaM fluorescence profile is not skewed by the ATP:ADP ratio or lactate biosensors because the excitation profiles do not overlap.

**Figure 2.**
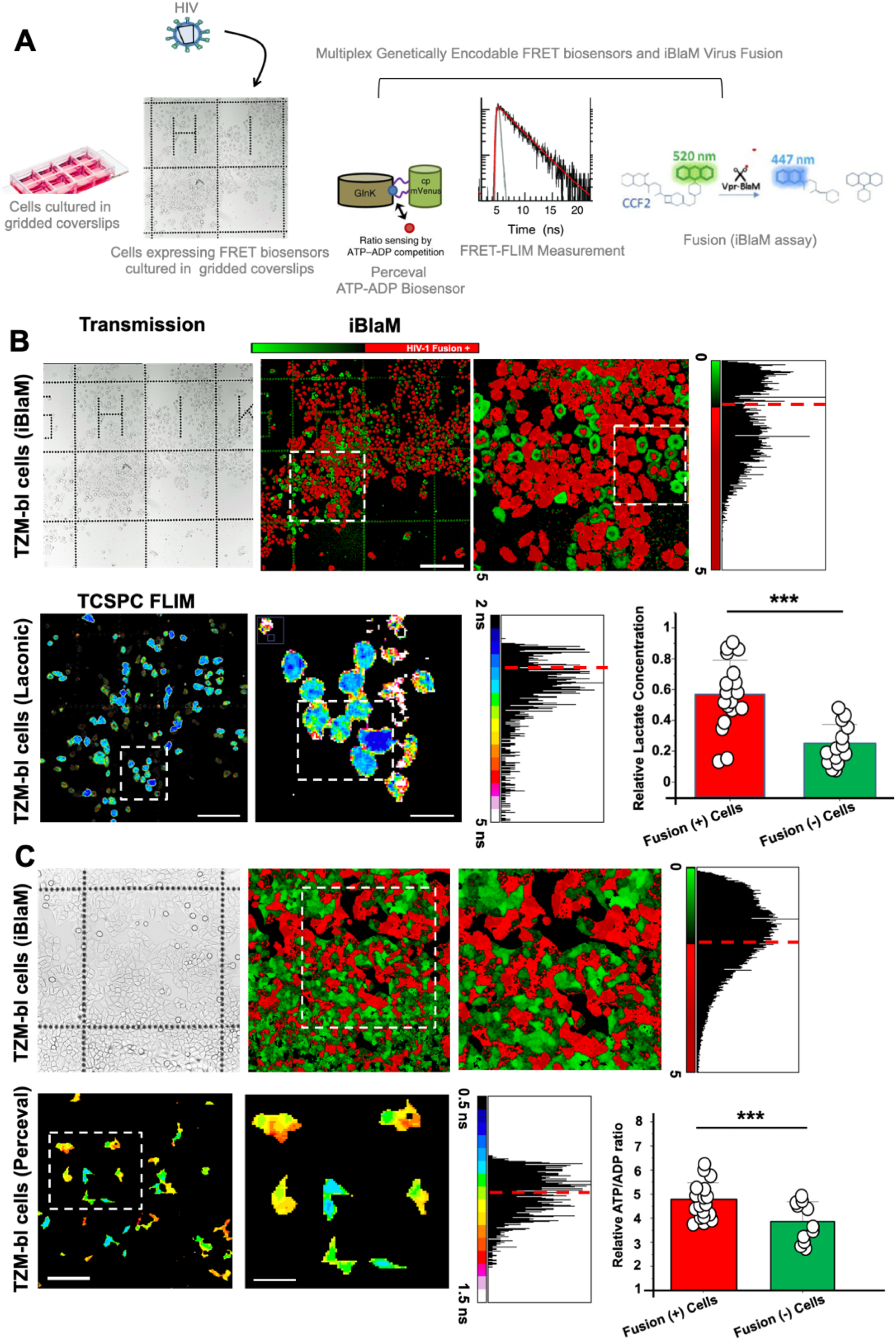
Relative lactate and ATP/ADP concentrations in single cells correlate with HIV-1 fusion. A.) Approach to evaluate if basal, single-cell ATP:ADP ratios or intracellular Lactate concentrations could predict HIV-1_JR-FL_ fusion. TZM-bl cells were seeded onto an Ibidi® 8-well gridded dish and transiently transfected with metabolic sensing biosensors Laconic or Perceval. FLIM images of cells transiently expressing these biosensors were recorded, denoting the location of the cells on the grid. These cells were then treated with HIV-1_JR-FL,_ and after 90 minutes, viruses were washed away and a β-lactamase assay was recorded on the same cells as a readout of HIV-1_JR-FL_ fusion. B.) Representative images of CCF2-loaded cells in Ibidi® 8-well gridded dishes recorded 90 minutes after HIV-1_JR-FL_ infection (top row) and basal FLIM images of intracellular lactate biosensor Laconic-expressing TZM-bl cells taken before HIV-1_JR-FL_ treatment (bottom row) illustrating cells with higher intracellular lactate concentrations (warm colours) were more likely to be scored fusion-positive for HIV-1_JR-FL_ fusion by the β-lactamase assay (bottom row). C.) Representative images of CCF2-loaded cells in Ibidi® 8-well gridded dishes recorded 90 minutes after HIV-1_JR-FL_ treatment (top row) and basal FLIM images of intracellular ATP:ADP biosensor Perceval-expressing TZM-bl cells taken before HIV-1_JR-FL_ treatment (bottom row) illustrating cells with higher intracellular ATP:ADP concentrations (cool colours) were more likely to be scored fusion-positive for HIV-1_JR-FL_ fusion by the β-lactamase assay (bottom row). *** p<0.001 as determined by Student T test of three independent experiments.

Interestingly, single cells with a higher relative lactate concentration were more likely to be fusion positive as determined by the BlaM assay than cells with a lower relative lactate concentration (Figure 2B). Similarly, cells with a reportedly higher ATP:ADP ratio were more likely to be fusion positive than cells with a lower relative ATP:ADP ratio (Figure 2C). Therefore, our data suggests that cells with a higher glycolytic flux are more likely to be infected by HIV-1_JR-FL_, and this bias for glycolytically-active cells occurs at the point of HIV-1 fusion.

### Inhibition of Glycolysis Blocks HIV-1 Fusion at Hemifusion

We tested whether acute arrest of glycolysis via incubation with increasing concentrations of 2-DG would inhibit HIV-1_JR-FL_ fusion as reported by the BLaM assay. Treatment of cells with increasing concentrations of 2-DG lead to a reduction in both the ATP:ADP ratio and lactate levels in biosensor-expressing cells with minimal loss in cell viability as determined by propidium iodine staining (Supplemental Figure 1, Supplemental Figure 3).

Pre-treatment of TZM-bl cells with increasing concentrations of 2-DG resulted in dose-dependent decreases in the relative amounts of lactate and ATP:ADP and a concomitant inhibition of HIV-1_JR-FL_ fusion (Figure 3A and 3B). Acute treatment with 100mM 2-DG blocked HIV-1_JR-FL_ fusion by nearly 80%. We similarly observed that increasing concentrations of 2-DG disrupted fusion of VSV-G-pseudotyped HIV-1 fusion, albeit to a lesser extent. Glucose deprivation resulted in similar reductions in virus-cell fusion events (Supplemental Figure 3B). Furthermore, acute treatment of TZM-bl cells with 2-DG did not lead to any visible alterations in virus attachment or surface expression of CD4 receptor or CCR5 co-receptor (Supplemental Figure 3C). Therefore, glycolysis appears to be a major regulator of successful HIV-1 fusion in human target cells.

**Figure 3.**
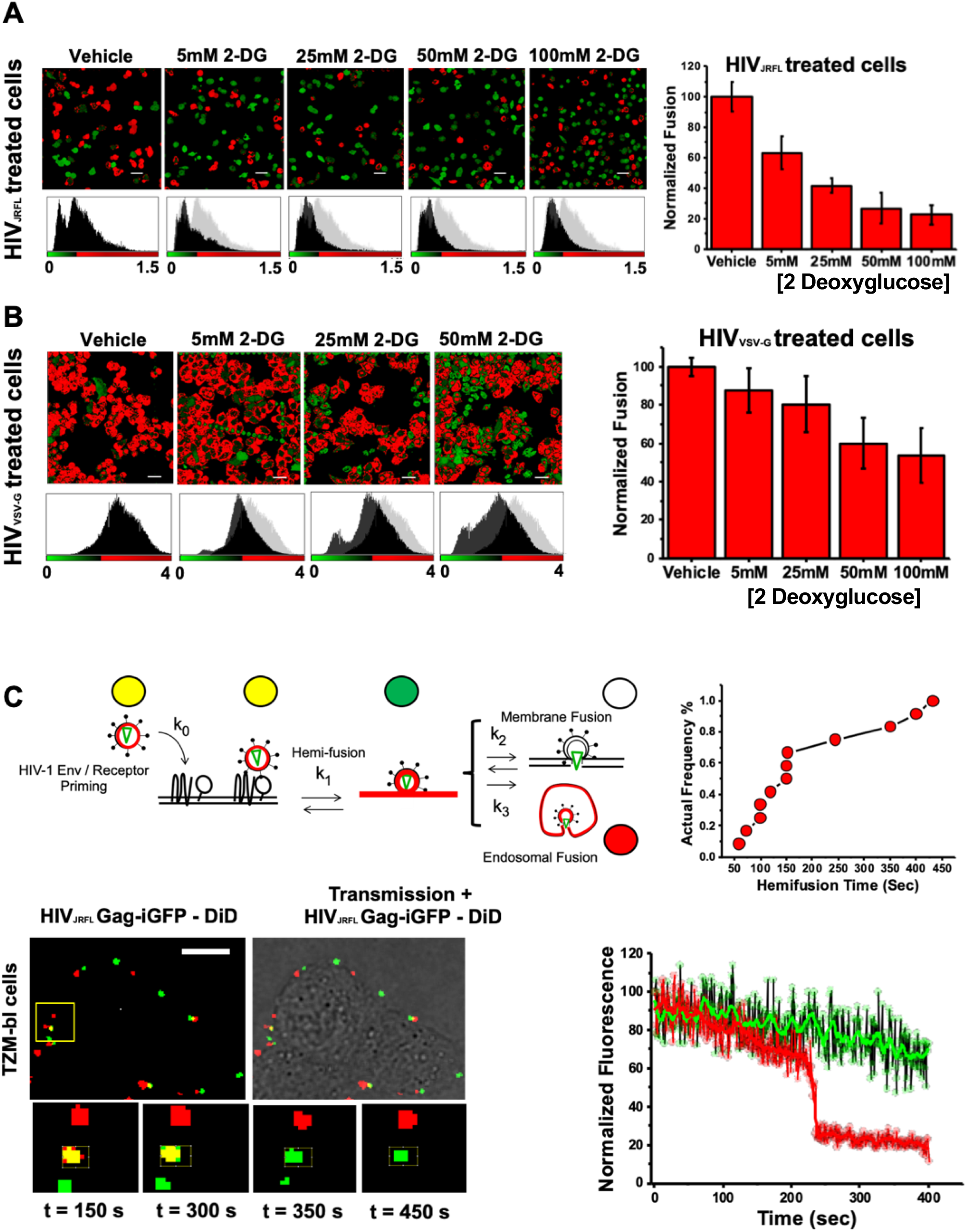
Addition of 2DG arrests HIV-1 fusion at the hemifusion stage. A.) As determined by β-lactamase assay and normalised to vehicle-treated control, increasing concentrations of 2-DG led to reductions in viral fusion for HIV-1_JR-FL._ (mean of three independent experiments). B.) As determined by β-lactamase assay and normalised to vehicle-treated control, increasing concentrations of 2-DG led to reductions in viral fusion for HIV-1_VSV-G_ (mean of three independent experiments). C.) Cartoon diagram illustrating the concept of single-particle tracking with double-labelled virions with DiD and eGFP-gag. Briefly, double-labelled virions entering via endocytosis will have their eGFP-gag signal infinitely diluted during endosomal fusion whilst DiD signal is retained in the endosome, that is mobile. Virions entering via plasma membrane fusion will have their DiD signal infinitely diliuted in the plasma membrane whereas the eGFP-gag signal is retained and mobile. Hemifusion is denoted when DiD signal infinitely diliuted in the plasma membrane whereas the eGFP-gag signal is retained and immobile Kinetics of the individual hemifusion events plotted as cumulative distributions as a function of time (top row, right). In the bottom row, left, there is a representative panel of images illustrating doubled-labelled HIV-1_JRFL_ particles losing DiD signal (red) and maintaining immobile eGFP signal (green) when attempting fusion in 2-DG treated cells, suggesting arrest at hemifusion (n=15). In the bottom row, right, there are representative panel of images illustrating doubled-labelled HIV-1_JRFL_ particles losing eGFP signal and maintaining DiD signal when fusing with untreated cells, suggesting normal endocytosis (n=15). *p<0.5, **p<0.01 *** p<0.001 as determined by one-way ANOVA.

To further characterise how glycolytic inhibition abrogated HIV-1 fusion in TZM-bl cells, we tracked single virus particles upon exposure to cells. To accomplish this, we co-labelled particles with the lipophilic dye DiD, which incorporates into the viral lipid bilayer, and eGFP fused to Gag to identify the virus core^28^. DiD loss of a particle marked by eGFP-Gag signals lipid mixing between viral and cellular membranes. Furthermore, differences between the surface area of the plasma membrane and endosomal membranes result in differential dilution of the lipophilic dye. As such, lipid mixing at the plasma membrane results in complete loss of the dye, whereas mixing at endosomal membranes has relatively little impact on the lipophilic dye (Figure 3C). Irrespective of the initial site of lipid mixing, loss of eGFP-Gag signals that pore formation is complete and cytosolic disintegration of viral core has occurred.

Upon infection of untreated TZM-bl cells, punctate structures of Gag-eGFP and the lipophilic dye DiD co-localize together at time = 0 sec turn red (t_1/2_ = 10.25 minutes). This confirms previous studies that HIV-1_JR-FL_ particles enter TZM-bl cells via endocytosis as Gag-GFP signal is lost indicating infinite dilution into the host cytosol whilst DiD signal (red) persists as it remains in the endosomal membrane (Supplemental Figure 3D) (Miyauchi et al. 2009; Jones and Padilla-Parra 2015). In contrast, when tracking yellow particles on TZM-bl cells treated with 2DG (100 mM) they turned green (Figure 3C) indicating DiD dilution at the plasma membrane in the absence of viral content release (Gag-GFP). This process is suggestive of the hemifusion intermediate (Miyauchi et al. 2009; Vega et al. 2011; Markosyan et al. 2005). Moreover, the kinetics of lipid mixing at the plasma membrane corroborates previously reported analyses of hemifusion events (Vega et al. 2011). Specifically, upon infection of 2-DG-treated TZM-bl cells, punctate structures of Gag-eGFP and the lipophilic dye DiD co-localize together at time = 0 sec turn green with a t_1/2_ = 2.53 minutes). Interestingly, this analysis uncovered two distinct types of DiD disappearance events. One population exhibited a nearly-asymptotic loss of DiD fluorescence (Figure 3C), whereas another population had a slow DiD fluorescence decay lasting up to 10 minutes or longer (Supplemental Figure 3E). We observed that 7% of single virions rapidly released their lipophilic marker into the plasma membrane, which is similar to previously reported SVT analyses of these events (Vega et al. 2011). In summary, we have shown that cells treated with increasing concentrations of 2-DG increasingly reduce the amount of fused virus particles in reporter TZM-bl cell lines and that this arrest to fusion by JR-FL pseudoviruses occurred at the hemifusion stage.

### Single-Cell Metabolic State is linked to Plasma Membrane Cholesterol Content

Several studies have linked the inhibition of a variety of metabolic pathways to the availability of lipids in the plasma membrane (Hao et al. 2002; Eid et al. 2017; Habbeger et al. 2012), several of which are important for successful virus entry, particularly cholesterol (Campbell et al. 2001; Yang et al. 2017; Rappocciolo et al. 2014; Delucia and Rinaldo 2018). To assess whether the inhibition of glycolysis was tied to the amount of plasma membrane cholesterol, we stained for cholesterol with filipin after treating cells acutely with increasing concentrations with 2-DG. Filipin has been shown to report plasma membrane cholesterol content without significantly disrupting the architecture of the plasma membrane (Wilhelm et al. 2019). Indeed, in both TZM-bl and MT4 cells, we noticed that acute treatment with 2-DG led to reductions in plasma membrane cholesterol by 21% and nearly 50% at 100mM 2-DG concentration in T cells and reporter cells, respectively (Figure 4A). Immunofluorescence imaging further confirmed these results, illustrating substantial loss of cholesterol from the plasma membrane surface (Figure 4B). Furthermore, treatment with methyl-β-cyclodextrin (MBCD), an acute cholesterol depletion reagent that strictly acts at the cell surface (Zidovetzki and Levitan 2008), led to similar reductions in cholesterol staining signal as 2-DG (Supplemental Figure 4A). Interestingly, treatment with increasing concentrations of ATP synthase inhibitor oligomycin did not lead to a significant decrease in plasma membrane cholesterol (Supplemental Figure 4B), and these results were confirmed by immunofluorescence labelling with filipin (Supplemental Figure 4C).

**Figure 4.**
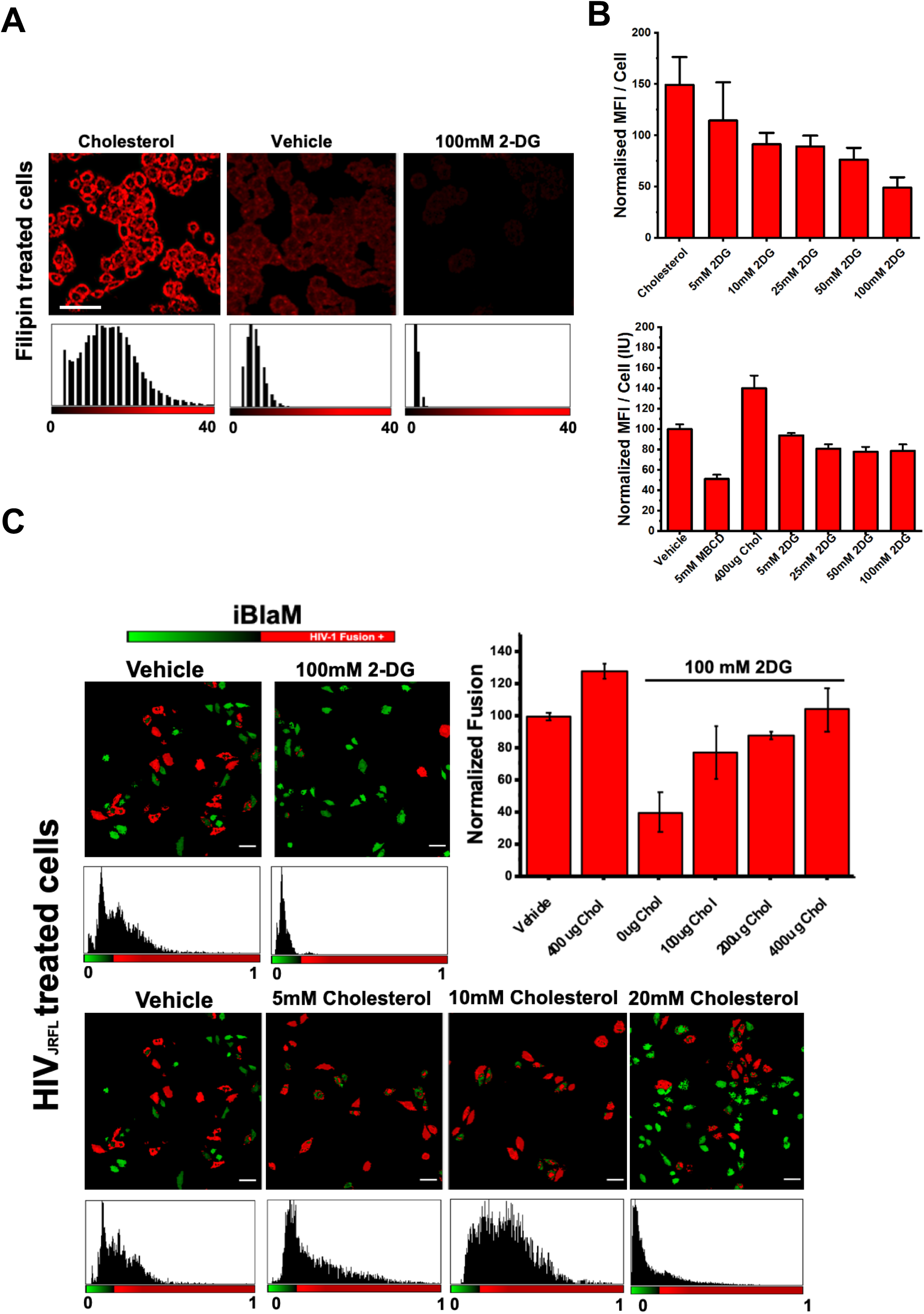
Addition of 2DG sequesters cholesterol from the cell membrane. A.) Representative images depicting filipin mean fluorescence intensity per cell indicating 2 hour incubation of increasing concentrations of 2-DG decreases surface cholesterol in TZM-bl cells. B.) Bar charts depicting mean fluorescence intensity per cell normalised to vehicle in TZM-bl cells (top) and MT4 cells (bottom) in cells treated with 400µg cholesterol, 5mM MBCD and increasing concentrations of 2-DG. C.) Representative images (left) and bar chart (right) depicting percent of fusion positive cells relative to vehicle control illustrating increasing concentrations of cholesterol rescues fusion in 2-DG treated cells. Bar charts depicting filipin mean fluorescence intensity per T cell normalised to vehicle, indicating 2 hour incubation of increasing concentrations of 2-DG, but not oligomycin decreases surface cholesterol. *p<0.5, **p<0.01 *** p<0.001 as determined by one-way ANOVA.

To determine if the loss of plasma membrane cholesterol via the inhibition of glycolysis was responsible for the block in virus-cell fusion, we pre-treated our reporter cells with 100mM 2-DG and with increasing concentrations of water soluble cholesterol to determine if restoration of plasma membrane cholesterol in the face of glycolytic inhibition would restore HIV-1_JR-FL_ fusion in reporter cells. We observed a near-complete rescue of virus fusion when cells were supplemented with 100µg/mL water-soluble cholesterol as reported by the β-lactamase assay (Figure 4C). Our results were confirmed by confocal microscopy of β-lactamase-treated cells (Figure 4D) Therefore, it appears that inhibition of glycolysis leads to loss of plasma membrane cholesterol, which negatively impacts HIV-1_JR-FL_ fusion.

### Single-Cell Glycolytic Activity Influences HIV-1 Fusion by Affecting Membrane Order and Tension

To assess whether inhibition of glycolytic activity alters the membrane order (i.e. a measurement of membrane packing) in single HIV-1 reporter cells, we utilised a recently established sensor for membrane order, FliptR (Colom et al. 2018). FliptR, a planarizable push-pull probe with high photostability (Colom et al. 2018; Soleimanpour et al. 2016) constructed from two large dithienothiophene flippers, segregates with similar efficiency into different membrane phases (Molin et al. 2015) and maintains the cell’s innate membrane order. We subsequently loaded our cells with FliptR and determined the effects of acute glycolysis inhibition on membrane order in TZM-bl cells and compared these effects to cells supplemented with or depleted of cholesterol (Supplemental Figure 5A). The distribution of lifetimes was homogenous throughout the entirety of the cell regardless of cholesterol enrichment or depletion (Supplemental Figure 5C), indicating that the liquid-ordered and disordered phases are mixed in our spatiotemporal resolution, which has been reported previously in similar TCSPC platforms (Colom et al. 2018; Kilin et al. 2015). Treatment with MBCD showed reductions in lifetime by over 200ps at 1mM concentrations (Figure 5a, 5b), indicating a drop in membrane order. Interestingly, cells treated with 2-DG showed similar reductions in membrane order when compared with control cells, supporting that acute 2-DG treatment leads to a loss in high order lipids in the plasma membrane (Supplemental Figure 5A). These results were confirmed by comparisons of extracted TCSPC histograms (Supplemental Figure 5D). Interestingly, supplementation of 2-DG treated cells with water-soluble cholesterol capable of plasma membrane insertion restored membrane order lifetime values to vehicle-treated values. Notably, increasing membrane tension by subjection to hypo-osmotic shock did not lead to a significant change in the membrane order of the cell.

**Figure 5.**
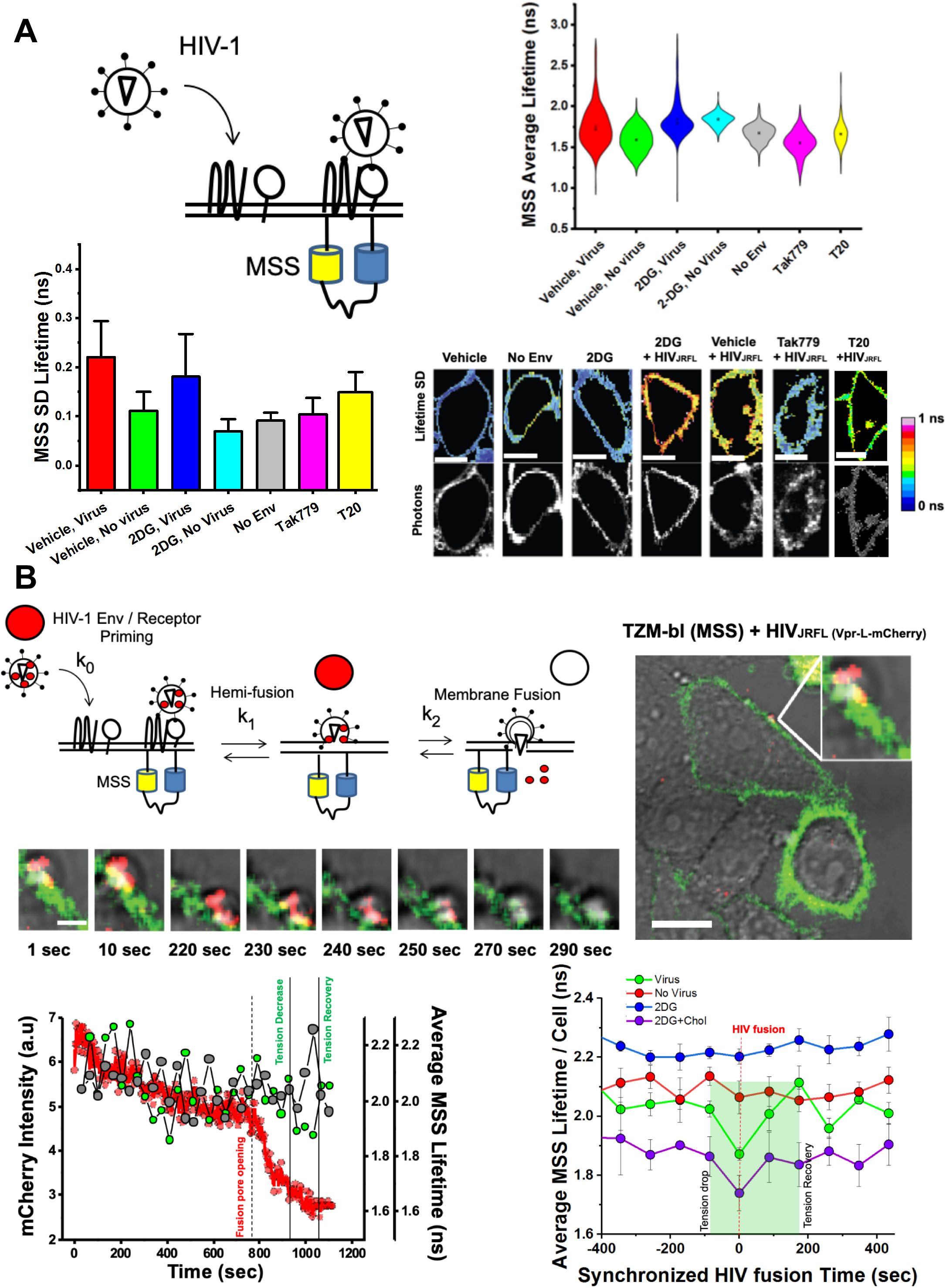
Multiplexed FLIM with SVT reveals a drop in local tension during single HIV-1 fusion in live cells. A.) (Top Left) Cartoon diagram depicting approach to determine plasma membrane tension fluctuations during virus entry. (Top Right) Violin plots of average MSS lifetimes per cell per frame during FLIM acquisition during viral entry. (Bottom Left) Standard deviation of MSS lifetimes per cell during real-time FLIM acquisition during viral entry. (Bottom Right) Representative images of the standard deviation of MSS lifetimes per cell for each condition during viral entry.* p<0.05 ** p<0.01 *** p<0.001 as determined by one-way ANOVA of three independent experiments. B.) (Top Left) Approach to combine SPT and MSS lifetime values to analyse local membrane tension during HIV-1_JR-FL_ entry. Representative images of a single Vpr-mCherry JR-FL pseudotyped VLP entering MSS-transfected TZM-bl cells are also depicted. (Bottom Left) Line graphs of local lifetimes in overlapping Vpr-mCherry regions (green) and non-overlapping regions (grey) for a single virus particle during entry (loss of mCherry signal, red). (Bottom Right) Compiled local MSS lifetimes during multiple viral fusion events (green) and respective virus-free regions within the same cell (red) compared to regions of 2-DG treated cells where viruses were incapable of fusion (blue) or rescued events due to cholesterol treatment (purple). *p<0.5, **p<0.01 *** p<0.001 as determined by one-way ANOVA.

We utilised a previously published FRET-based reporter of membrane tension, MSS (Li et al. 2018), to determine changes in membrane-tension of single cells during acute glycolytic inhibition. MSS is an ECFP-YPet FRET-based membrane-bound tension reporter constructed from an elastic protein tension sensing module and two anchoring proteins linked with lipid molecules in raft and non-raft regions of the plasma membrane (Li et al. 2018). Upon treatment with identical conditions as the FliptR reporter, we detected a similar pattern in changes in CFP lifetime changes (Supplemental Figure 5E) and FRET efficiency (ECFP/YPet ratio) (Supplemental Figure 5F) when compared to vehicle-treated cells. Acute cholesterol depletion with mild concentrations of MBCD drastically increased the donor lifetime over 100ps (i.e. Δτ=101ps) (Supplemental Figure 5E), indicating that cholesterol is a requirement to buffer plasma membrane tension in cells. Remarkably, 2-DG treatment led to even larger increases in donor lifetime (i.e. Δτ=165ps), indicating a reduction in FRET efficiency and an increase distance (i.e. stretch and tension) between donor and acceptor fluorophores (Supplemental Figure 5E). Furthermore, supplementing 2-DG treated cells with water-soluble cholesterol restored membrane tension values to control levels.

Cellular plasma membrane tension is a major regulator of several cellular processes, including those that favour or disfavor virus entry (Yang et al. 2015, 2016, 2017; Kozlov and Chernomordik 2015). Since we observed a connection between the glycolytic state of the cell and its corresponding membrane order and tension, we sought to measure how HIV-1_JR-FL_ impacts membrane tension during the fusion process. To accomplish this, we analysed single cells transiently expressing the MSS tension reporter during mCherry-Gag labelled HIV-1_JR-FL_ fusion via single particle tracking concomitant with FLIM of the MSS reporter (Figure 5A). Virus fusion was determined by the rapid loss of mCherry signal. Interestingly, we initially observed that cells treated with HIV-1_JR-FL_ possessed larger fluctuations in CFP donor lifetimes regardless of 2-DG treatment, indicating that cells exposed to virus particles actively undergo changes to membrane tension in response to HIV-1_JR-FL_ (Figure 6A, top right panel). These broad fluctuations in membrane tension were Env-dependent, as non-enveloped HIV-1 particles failed to trigger them during SVT acquisition. Pre-treatment of cells with CCR5 HIV-1 entry inhibitors Tak779, a CCR5 antagonist which inhibits co-receptor engagement by HIV-1 gp120, and T20, an HIV-1 fusion inhibitor which competitively binds to gp41 and blocks its post-fusion structure suggested that these changes were CCR5 dependent. Indeed, MSS CFP-donor lifetime fluctuations in Tak779-treated cells were narrowly similar to non-enveloped, naked HIV-1 virions (Figure 6B). In contrast, upon T20 treatment, these CFP donor lifetime fluctuations were partially restored, confirming that fluctuations in whole-cell target membrane tension during HIV-1_JR-FL_ entry might be CCR5-dependent. These results were confirmed by analysing the pixel-by-pixel lifetime standard deviations of MSS-containing regions of whole cells during virus entry with and without entry or metabolic inhibitor addition (Figure 6A, bottom two panels).

**Figure 6.**
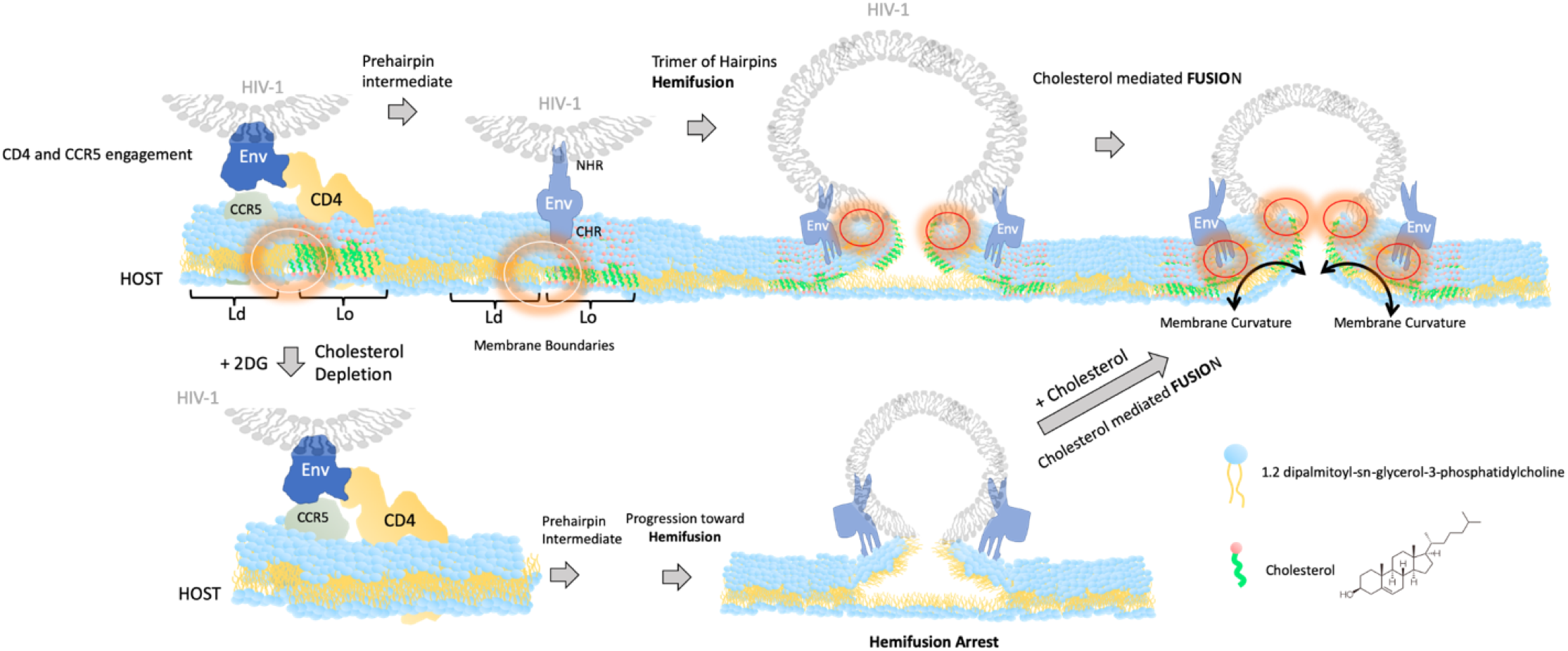
Cholesterol availability regulates the transition between HIV-1 hemifusion and fusion. HIV-1 Env sequentially interacts with CD4 and CCR5 at the boundaries of ordered lipid domains. The progression toward hemifusion, however, is not cholesterol dependent as cells treated with 2DG showed a reduced concentration of cholesterol and were able to progress toward HIV-1 hemifusion. Viruses exposed to cells pre-treated with 2DG were arrested right at hemifusion and only the addition of cholesterol rescued full fusion suggesting that cholesterol is a limiting factor in the HIV-1 fusion reaction in the process of fusion pore formation. White circles (left panels) represent lipid boundaries where cholesterol might induce HIV-1 Env priming and favor the prehairpin intermediate. Red circles (right panels) represent a potential role of cholesterol during fusion pore formation and membrane curvature.

We also evaluated local changes in membrane tension during HIV-1_JR-FL_ entry into individual cells. We tracked the MSS CFP donor lifetime in 3 x 3 pixel regions at the plasma membrane overlapping with mCherry-Gag labelled HIV-1_JR-FL_ during virus approach by multiplexing our FLIM with SVT. Virus fusion was determined by the rapid loss of mCherry-Gag signal. In reporter cells free from any treatment, we noticed a sharp drop in MSS donor lifetimes in plasma membrane regions coinciding with the disappearance of proximal mCherry-Gag signal (Figure 6B, bottom left panel, green curve). In comparison, plasma membrane regions lacking virus particles in the same cell did not display any sharp decreases in MSS donor lifetime (Figure 6B, bottom left panel, grey curve). This local drop in MSS lifetime (i.e. membrane tension) overlapping with mCherry-Gag signal disappearance in space and time was statistically significant when compared to mCherry-Gag signal-free regions of similar size in the same cell (Supplemental Figure 6B). However, there was no statistically significant difference in baseline plasma membrane tension values between these two regions at baseline, before loss of mCherry-Gag signal (Figure 6B, bottom right panel, Supplemental Figure 6C). Furthermore, this local drop in MSS CFP donor lifetime was transient, as there was rapid restoration of the MSS CFP donor lifetime almost immediately after the loss of the mCherry-Gag signal (Figure 6B, bottom right panel). In fact, of the total events analysed (n=10), there was a statistically significant rebound in MSS CFP donor lifetime values of plasma membrane regions near mCherry-Gag signal that was recently departed compared to baseline MSS lifetime value (Figure 6B, bottom right panel, Supplemental Figure 6D). These results indicate that during HIV-1_JR-FL_ entry in reporter cells there is a local reduction in plasma membrane tension that is immediately restored following virus fusion.

When analysing MSS CFP donor lifetimes in 3 x 3 pixel plasma membrane regions overlapping with mCherry-Gag-containing viruses attempting to enter cells treated with 2-DG, we noticed that viruses unable to enter cells were associated with plasma membrane regions with relatively high membrane tension values when compared to vehicle-treated cells (Figure 6B, bottom right panel). These higher plasma membrane tension values were also present when analysing the whole cell (Supplemental Figure 6A). Additionally, we failed to find drops in MSS lifetime values during the loss of mCherry-Gag signal (Figure 6B, bottom right panel) when compared to local PM region values in all viruses analysed (n=10), indicating that the localised nature of the plasma membrane tension drop during viral entry is inhibited by 2-DG. Remarkably, we recovered MSS lifetime profile traces similar to non-treated cells when 2-DGtreated cells were supplemented with water-soluble cholesterol (Figure 6B, bottom right panel). Altogether, this shows that a cell’s glycolytic activity sets surface cholesterol levels and plasma membrane tension that determines success of HIV-1 fusion and thus ultimately HIV-1 infection.

## DISCUSSION

Recent studies have stressed the impact of host metabolism on HIV infection (Buck et al. 2017; Pallett et al. 2019; Valle-Casuso et al. 2018). It has been shown that differences in HIV-1 susceptibility between naive and more differentiated T cells were related to metabolic activity, in particular to oxidative phosphorylation and glycolysis (Valle-Casuso et al. 2018). Inhibition of glycolysis impaired HIV-1 infection in cell culture in all CD4+ T cell types. The mechanism behind these results and the link between metabolism and HIV-1 fusion, however has remained elusive until now. Our data with T cells utilising HIV-1 pseudoparticles corroborates the finding that acute inhibition of glycolysis inhibits productive infection (Figure 1). Moreover, we also show that inhibition of glycolysis specifically sequesters cholesterol from the plasma membrane in both T cells and TZM-bl cells (Figure 4). The combination of genetically encoded calibrated FRET-based biosensors detecting metabolic activity permitted the use single-cell ATP/ADP and lactate as an indirect reporter for glycolytic activity (Figure 1 and 2). These experiments were combined with single cell fusion and infection assays and revealed two important findings. First, the metabolic landscape predetermined a cell’s propensity for HIV-1 infection.

Various envelopes were utilised to pseudotype our HIV-1 particles: VSV-G, NL4.3 and JR-FL. It is well-established that VSV-G pseudotyped virions enter host cells via endocytosis (Sun et al. 2005). However, the entry site of HIV-1 envelopes JRFL and NL4.3 are debated and argued to be cell-type dependent (Melikyan 2015). It is thought that HIV-1 R5- and X4-tropic virions enter both primary T cells and T cell lines specifically at the plasma membrane, whereas endocytosis may be dispensable (Herold et al. 2014; Kondo et al. 2015;). However, in reporter cells, it has been put forth that HIV-1 may productively enter host cells via endosomes (Vega et al. 2011). Our report illustrates that glycolytic activity and its regulation of physical properties in the plasma membrane via cholesterol content are universally required regardless of the entry pathway. Curiously, we detected that this dependence of glycolytic activity and virus entry seemed more relevant to HIV-1 JRFL and NL4.3 as opposed to VSV-G, and the entry path may reflect these differences, as cholesterol content between the plasma membrane and endosomes are often quite different.

Plasma membrane lipid bilayers contain a variety of lipids which are thought to be laterally segregated into domains to regulate fusion, endocytosis and signal transduction (Jacobson et al. 2007; Pike 2009). Lo domains are thought to be dominated by saturated lipids (i.e. sphingolipids) and sterols (i.e. cholesterol) surrounded by a pool of unsaturated phospholipids constituting Ld domains. The contributions of cholesterol and the boundaries between Lo and Ld domains to the successful entry of viruses including HIV-1 is well-established. Membrane order and tension are significant contributors to several membrane-regulated processes in human cells important for viral infection, including viral entry (i.e. fusion and endocytosis) and egress (Ewers and Helenius 2011; Kozlov and Chernomordik 2015). Recently, several studies have offered a link between metabolic activity and sensing in live cells to cellular membrane tension indicating that various membrane-regulated processes are controlled by master metabolic sentinels (Rispal et al. 2015; Bourgoint et al. 2018; Riggi et al. 2018). Our results further build upon this link suggesting that high glycolytic activity may increase susceptibility of cells to support HIV-1 entry due to membrane tension regulation.

Specifically, our observation that decreased virus entry occurs following glycolytic inhibition could be linked to alterations in either membrane order or tension. We hypothesise that glycolysis directly, through the production of acetyl-CoA precursors, or indirectly regulates cholesterol in cellular membranes as the addition of 2-DG inhibited cholesterol accumulation without affecting CD4 or co-receptor expression levels (Supplementary Figure 3). This loss in cholesterol led to drastic increases in membrane tension and loss of membrane order, owing to the sequestration of cholesterol from the plasma membrane (Figure 5 and Supplementary Figure 5). These two aspects are crucial for the HIV-1 fusion reaction for many reasons. First, it has been shown that lipid-ordered (i.e. cholesterol-laden) and lipid-disordered membrane domains (also termed raft boundaries) can act as an attractor for HIV Env (Molotkovsky et al. 2018; Yang et al. 2017). Both experimental and theoretical studies show that the presence of raft boundaries in the vicinity of HIV-1 entry sites reduces the energy barrier required to perform virus-cell fusion (Yang et al. 2016). Moreover, the line tension created at lipid domain boundaries drive gp41 mediated fusion, owing to the mismatches in lipid length occurring at Lo and Ld boundaries, most likely creating membrane curvature which favours virion fusion (Yang et al. 2016)) (Figure 6). Cholesterol and other ordered lipids are known to maintain negative spontaneous curved membrane domains, which facilitates the creation of a fusion stalk by reducing its formation energy in addition to enabling its progression to fusion pore formation (Meher and Chakraborty 2019). The sequestration of cholesterol from the host membrane in glycolytically-inactivated cells arrests HIV fusion, as this line-tension necessary to drive fusion is diminished. Furthermore, sequestration of cholesterol from the host plasma membrane ostensibly increases membrane tension (Figure 6), which has been illustrated previously to collapse hemifusion intermediates (Floyd et al. 2008) in addition to preventing the formation of highly curved intermediates, disfavouring packing defect formation and the generation of nucleation points for two membranes to be joined (Meher and Chakraborty 2019). Moreover, this increase in membrane tension might in turn have an effect on the optimal hydrophobic equilibrium between ordered and disordered lipid domains needed to complete the fusion reaction (Yang et al. 2016, 2017) and (Figure 5).

We hypothesise that there is a range of hydrophobic mismatch between ordered and disordered lipid domains that renders HIV fusion plausible. When this equilibrium is lost (both toward high or low tension) the energy required for gp41 fusion peptide to complete fusion (expressed in tens of kBT, (Yang et al. 2016)) is not compatible with the completion of the fusion reaction. This is why depletion of cholesterol might allow HIV Env to be primed with CD4 and CCR5 followed by arrest at hemifusion (Figure 3) even if overall tension in the host cell membrane is increased (Figure 6). Moreover, our data suggests that cholesterol is a limiting factor for the transition between HIV-1 hemifusion and fusion (Figure 6) and not necessarily during the sequential interactions between HIV-1 Env with CD4 and CCR5 co-receptor at phase boundaries (Yang et al. 2017). Indeed, in cells treated with 2DG where cholesterol is sequestered, HIV-1 was able to progress until hemifusion. The external addition of cholesterol allowed full fusion to be completed (Figure 6).

Our results indicate that glycolytic activity is needed to maintain higher lipid order and lower levels of membrane tension inside cells, two characteristics that promote HIV-1 fusion with cells. Moreover, we have also shown that HIV-1 induces broad fluctuations in host global membrane tension during virus entry, yet require localised, transient reductions in plasma membrane tension in order to enter their host cell. In brief, our work shows that glycolytic inhibition with 2-DG creates changes in global and local membrane tension that restricts HIV-1 fusion at the hemifusion stage and that this inhibition can be tied to cholesterol availability. Importantly, as glycolytic activity sets membrane tension levels by altering cholesterol levels of the cell, therapeutic strategies that specifically target membrane cholesterol may find utility in HIV-1-infected individuals or in those at increased risk of HIV-1 infection.

## MATERIAL AND METHODS

### Cell Culture

TZM-bl and Lenti-X-293T cells were cultured in either complete Dulbecco’s Modified Eagle Medium (DMEM) or DMEM F-12 (Life Technologies), respectively, both of which contained 10% fetal bovine serum (FBS), 1% Penicillin-Streptomycin (PS) and 1% L-glutamine (LGlut). Cells were maintained at 37°C which provided 5% CO_2_. MT4 cells were cultured in RPMI containing 10% FBS, 1% PS and 1% L-Glut.

### Reagents and antibodies

All chemical and biochemical reagents were obtained from the following sources: The Flipper-TR probe was purchased from Spirochrome (Geneva, Switzerland). Antibodies against CD4 conjugated to PercP Cy5.5 (ab161a1) and CCR5 conjugated to FITC (ab11466) were from BioLegend (San Diego, CA, USA) and Abcam (Cambridge, UK), respectively. β-lactamase CCF2-AM loading solutions from the Live-BLAzer FRET B/G Loading Kit was obtained from Life Technologies (Carlsbad, CA, USA); Filipin, 2-deoxy-glucose (2-DG), Methyl-β-cyclodextrin (MBCD) and water-soluble cholesterol (Xu et al. 2010) was obtained from Sigma-Aldrich (St. Louis, MO, USA).

### Plasmids Transfections

The pR8ΔEnv plasmid (encoding HIV-1 genome harbouring a deletion within Env), pcRev, VPR-BlaM, Gag-eGFP and VSV-G were kindly provided by Greg Melikyan (Emory University). The plasmid encoding the JR-FL envelope protein was a kind gift from James Binley (Torrey Pines Institute for Molecular Studies). The MSS sensor was kindly provided from Bo Liu (Dalian University of Technology). Perceval and Laconic plasmids were obtained from Addgene. Transient transfections of Perceval, Laconic (Addgene plasmid numbers 21737 and 44238, respectively) and the MSS probe was performed according to the manufacturer’s protocol for GeneJuice® (Novagen). Transiently-transfected cells were analysed by FLIM 24-48 hours later.

### Virus production

Vpr-BlaM-containing, JR-FL and VSV-G pseudotyped viral particles were produced via transfection of 60-70% confluent Lenti-X-HEK-293T cells seeded in T175 flasks. Component DNA plasmids were transfected into Lenti-X-HEK-293T cells via GeneJuice® (Novagen) according to manufacturer’s protocol. Specifically, cells were transfected with 2□μg pR8ΔEnv, 1□μg pcREV, 3□μg of the appropriate viral envelope (either VSV-G, the CCR5-tropic HIV-1 strain JR-FL, or the CXCR4-tropic HIV-1 strain NL4.3), and either 2□μg Vpr-BlaM if to be used for β-lactamase assays, or 3μg of eGFP-GagΔEnv if being used for flow cytometry experiments. Alternatively, if to be utilised for single particle tracking, 2□μg pR8ΔEnv, 3□μg Gag-eGFP-Δenv, 1□μg pcREV and 3□μg of the appropriate viral envelope were transfected into HEK-293T cells. If to be utilised for single particle tracking whilst recording MSS FLIM lifetime acquisition, 2□μg pR8ΔEnv, 2□μg mCherry-2xCL-YFP-Vpr, 1□μg pcREV and 3□μg of the appropriate viral envelope. All transfection mixtures were then added to cells supplemented in complete DMEM F12, upon which time they were incubated in a 37□°C, 5% CO_2_ incubator. 12□hours posttransfection, the medium was replaced with fresh, phenol-red free, complete DMEM F12 after washing with PBS. Cells were subsequently incubated for a further 24□hrs. If viruses were to be utilised for single particle tracking, 12 hours post-transfection, the transfection mixture was removed and cells were washed twice with PBS before being incubated at 37º C with DMEM F12 medium containing 10µM DiD (Life Technologies) for 4 hour. The staining mixture was subsequently removed, upon which time the cells were washed twice with PBS and DMEM F12 was then added again and incubated for an additional 24 hours prior to harvesting as follows. 48□hrs post-transfection for either Vpr-BlaM, double-labelled or immature virions, the supernatant containing virus particles was harvested, filtered with a 0.45□μm syringe filter (Sartorius Stedim Biotech), aliquoted and stored at −80□°C.

### Virus Titering

To titer the produced pseudovirions, 2 × 10^4^ TZM-bl cells were seeded in complete DMEM, in triplicate, in a 96-well plate (Falcon) and allowed to grow in a 37□°C, 5% CO_2_ incubator for 24 hours. Subsequently, the media was removed and replaced with 10-fold serially-dilutions of the produced pseudovirus particles. Aliquoted Vpr-BlaM-containing or double-labelled JR-FL and VSV-G pseudotyped viral particles additionally contained the β-galactosidase gene in the pr8ΔEnv construct. Cells were incubated for an additional 48 hours in a 37□°C, 5% CO_2_ incubator, upon which time the medium was removed, the cells were then washed with PBS and fixed with 2% paraformaldehyde for 10 minutes. After an additional PBS wash post-fixation, an X-gal (i.e. BCIG, or 5-bromo-4-chloro-3-indolyl-β-D-galactopyranoside) solution comprised of 500mM K_3_[Fe(CN)_6_], 250mM K_4_[Fe(CN)_6_], 1M MgCl_2_, PBS, and 50mg/mL Xgal was added with to the cells and incubated for two hours at 37□°C, 5% CO_2_ incubator in the dark. Afterwards, cells were washed with PBS and subsequently imaged with a 630nm excitation continuous laser (Leica, Manheim) while recording the emission spectra with a 650-750nm emission window, pixel by pixel (512 × 512) using a Leica SP8 X-SMD microscope with a lambda resolution of 12□nm, in order to count infection-positive cells for each 10-fold dilution to extrapolate the virus titer.

### β-lactamase Assay

Four-hours post-addition of the pseudotyped virus particles harbouring Vpr-BlaM at the specified MOI to the transfected TZM-bl cells, virus particles were removed, and cells were washed twice with PBS. After the washes, complete DMEM was added to each well and the cells were allowed to incubate at 37°C, 5% CO_2_. Cells were then loaded with CCF2-AM from the LiveBLAzer FRET—B/G Loading Kit (Life Technologies) and incubated at room temperature in the dark for 2□hrs. After this incubation step, the CCF2 was removed, and cells were washed with PBS and maintained in 2% FBS-containing, phenol-red free DMEM prior to imaging.

### β-lactamase assay spectral analysis

CCF2-loaded TZM-bl cells were excited with a 405nm continuous laser (Leica, Manheim) while recording the emission spectra with a 430-560nm window, pixel by pixel (512 x 512) using a Leica SP8 X-SMD microscope with a lambda resolution of 12□nm. Utilising ImageJ (http://imagej.nih.gov/ij/), the ratio of the blue (440-480nm, cleaved CCF2) to green (500-540nm, uncleaved CCF2) emission was calculated pixel by pixel using 20X objective. Statistical analyses of the BlaM data was performed using a two-tailed Fishers Exact Test (SigmaPlot, San Jose, CA). Cells were then removed of the CCF2 mixture and allowed to incubate for 24 hours in complete DMEM if they were to be analysed for infection by the B-Gal assay (below) for infection.

### β-galactosidase Assay

After imaging the CCF2-loaded TZM-bl cells, cells were washed with PBS and maintained in 2% FBS-containing, phenol-red free DMEM overnight for 12 hours in a 37□°C, 5% CO_2_ incubator. At least 36 hours after the addition of the pseudotyped virus particles, the medium was removed, and the cells were then washed with PBS and subsequently fixed with 2% paraformaldehyde for 10 minutes. After an additional PBS wash post-fixation, an X-gal (i.e. BCIG, or 5-bromo-4-chloro-3-indolyl-β-D-galactopyranoside) solution comprised of 500mM K_3_[Fe(CN)_6_], 250mM K_4_[Fe(CN)_6_], 1M MgCl_2_, PBS, and 50mg/mL Xgal was added to the cells, upon which time the cells were incubated for two hours in a 37□°C, 5% CO_2_ incubator in the dark. Afterwards, cells were washed with PBS and subsequently imaged with a 630nm excitation continuous laser (Leica, Manheim) while recording the emission spectra with a 650-750nm emission window, pixel by pixel (512 × 512) using a Leica SP8 X-SMD microscope with a lambda resolution of 12□nm, in order to identify infection-positive cells.

### Fluorescence lifetime imaging microscopy (FLIM)

Live TZM-bl cells expressing Perceval, Laconic or FlipTR were imaged using a SP8–X-SMD Leica microscope, whilst MSS-expressing cells were imaged using a SP8–X-SMD microscope with the FALCON module, both from Leica Microsystems (Manheim, Germany). Cells of interest were selected under either a 20x air-immersion or 63x oil-immersion objective (i.e. Perceval, Laconic and FlipTR) or a 100x/1.4 oil immersion objective (i.e. MSS). Perceval-expressing cells were excited using a 488□nm pulsed laser, Laconic-expressing cells were excited using a 470 nm pulsed laser, both tuned at 40□MHz coupled with single photon counting electronics (PicoHarp 300) and subsequently detected by hybrid external detectors in photon counting mode. FlipTR stained cells were excited with a 488 nm puled laser tuned at 20MHz. MSS-expressing cells were subjected to a 440nm picosecond pulsed diode laser PDL 800-B (PicoQuant) tuned at 40MHz in order to excite the FRET donor (CFP) while the emitted photons passing through the 450-480nm emission filter were detected using the internal hybrid detector in photon counting mode. Time-domain FLIM experiments were performed using a TCSPC approach operated by the FALCON module (Leica Microsystems, Manheim, Germany) integrated within the Leica SP8-X-SMD microscope. In order to remove artefacts caused by noise or photo–bleaching and insufficient signal to noise, cells with negligible amounts of bleaching and at least 250–1000 photons per pixel were only allowed in the analysis. Symphotime 64 software (Picoquant) was utilized to acquire the fluorescence decay of each pixel in individual cells expressing Perceval, Laconic, or Flipper-TR-stained cells, which was deconvoluted with the instrument response function (IRF) and fitted by a Marquandt nonlinear least–square algorithm with two–exponential models. The mean fluorescence lifetime (τ) was calculated as previously described using Symphotime. Statistical analysis of the lifetime data was performed using a two-tailed t-test (Origin, Northhampton, MA, USA).

### Flow Cytometry

Single-round infections were performed either with HIV-1_VSV-G_-eGFP-Gag or HIV-1_NL4.3_-eGFP-Gag. MT4 T cells were infected in triplicate (1 × 10^5^ cells/ well, 200uL) at an MOI of 1. Active HIV-1 infection was estimated by flow cytometry (BD LSRII, BD bioscience) as the percentage of eGFP-expressing, live MT4 T cells 48 hours after infection, and live cells were determined via LIVE/DEAD Fixable Near-IR Dead cell stain for 633/635 nm to stain dead cells following manufacturer’s instructions (Invitrogen).

### Extrapolating the ATP:ADP ratio and relative lactate concentrations and MSS Lifetimes

Utilizing Symphotime software (Picoquant Gmbh, Berlin, Germany), each whole cell expressing Perceval or Laconic was determined to be a region of interest. The region of interest with the smallest lifetime (τ_int_) was selected as the cell with the highest amount of ATP relative to ADP in the field of view analysed. From this region of interest, the short lifetime component (τ_1_) was fixed in all other analysed cells when acquiring the TCSPC fluorescence decay of each pixel, as this component possessed the highest amplitude *a*_1_, therefore representing the relative ATP concentration in the cell. The lifetime component τ_2_ was interpreted to represent the relative contribution from ADP binding to the Perceval sensor, and therefore its corresponding amplitude *a*_2_ was utilized to extract the relative ADP concentration. The resulting two-exponential decay which was deconvoluted with the instrument response function (IRF) and fitted by a Marquandt nonlinear least–square algorithm contained two amplitudes, *a*_1_ and *a*_2_. The ratio (i.e. *a*_1_/*a*_2_) of these two amplitudes was taken to generate the ATP:ADP ratio. To extract the relative lactate concentration from Laconic, a similar approach was utilised, where the amplitude *a*_1_ of the long lifetime component τ_1_ was divided by the sum of the amplitudes from the two-component exponential decay.

For the MSS probe, we utilised the internal FALCON module Leica Microsystems, Manheim, Germany) integrated within the Leica SP8-X-SMD microscope for our FLIM analysis. Whole cells expressing the MSS probe or 3 × 3 pixel-by-pixel regions of the MSS-labelled membrane overlapping with mCherry-labelled virions were chosen as our region of interest. To increase photon counts per pixel, we binned 5 × 5 pixels in addition to grouping frames of our acquisition in groups of 10. We utilised a three-exponential decay deconvoluted with the IRF and fitted by a Marquandt nonlinear least-square algorithm containing three amplitudes with the third long lifetime component fixed to the average of all cells analysed to isolate short τ_1_ and long τ_2_. The longer lifetime component (τ_2_) was chosen to represent the donor lifetime and therefore the membrane tension. When analysing 3 × 3 pixel-by-pixel regions of the MSS-labelled membrane overlapping with mCherry-labelled virions, to accommodate for low photon counting numbers, we utilised a single-exponential decay to extract MSS τ.

### Statistics

All statistical calculations (t-test, ANOVA, standard deviation and error) were calculated using Originlab software (Northhampton, USA)

## AUTHOR CONTRIBUTIONS

C.A.C., I.C-A. and M.I. performed infection and fusion experiments C.A.C. performed FLIM and SVT experiments. C.A.C. performed immunofluorescence and flow cytometry experiments. C.A.C. analysed the data from all experiments. C.A.C. and S.P-P. wrote the manuscript with comments from all authors. A.A.C., E.B.C. and S.P-P conceived the study. M.L.D., E.B.C., A.A.C., and S.P-P. co-supervised the study.

## ACKNOWLEDGEMENTS

We thank Luis Alvarez for technical support on advanced microscopy, acquisition, and analysis. We thanks Bo Liu for sharing the KMSS and MSS biosensors. C.A.C. acknowledges support from the Oxford-Cambridge Fellowship Program from the National Institutes of Health. S.P-P acknowledges funding from the Nuffield Department of Medicine Leadership Fellowship, Gobierno de Espana, Programa Estatal de I+d+I Orientada a los Retos de la Sociedad RTI2018-098415-A-I00 and all authors from the Wellcome Trust Core Award (203141). Work in the lab of A.A.C. is supported by the Intramural Research Program of the National Institutes of Health, National Cancer Institute, Center for Cancer Research. Wellcome Trust Principal Research Fellowship 100262Z/12/Z supported M.L.D. and European Commission Grant ERC-2014-AdG-670930 supported E.B.C.

The authors declare no competent interests.

---

**Figure 1 Supplement-1.** A.) (Left) Gating strategy to select and subsequently quantify single, live MT4 cells infected with either HIV-1_VSV-G_ or HIV-1_NL4.3_ harbouring eGFP-Gag during acute treatment with increasing concentrations of either 2-DG (pictured) or oligomycin. (Right) Bar charts depicting % eGFP-expressing cells as a marker of infection illustrating that acute treatment with 2-DG led to reductions in HIV-1_NL4.3_ infection in human MT4 T cells. B.) Dot plots representing lifetimes of ATP:ADP ratio biosensor Perceval extracted from live, single cells as regions of interest post-treatment for 2 hours with increasing oncentrations of 2-DG. C.) Representative intensity (left) and fluorescent lifetime imaging (right) of single cells transiently expressing ATP:ADP ratio sensor Perceval in single cells with increasing concentrations of glycolytic inhibitor 2-DG (scale bar, 50µm). D.) Dot plots representing lifetimes of intracellular lactate biosensor Laconic extracted from live, single cells as regions of interest post-treatment for 2 hours with increasing oncentrations of 2-DG. E.) Representative intensity (left) and fluorescent lifetime imaging (right) of single cells transiently expressing intracellular lactate sensor Laconic in single cells with increasing concentrations of glycolytic inhibitor 2-DG (scale bar, 50µm). F.) Bar charts representing the lifetime extracted from single cells expressing intracellular pH bionsensor pHRed indicating a lack of change in fluorescence lifetime. *p<0.5, **p<0.01 *** p<0.001 as determined by one-way student’s T-test.

**Figure 3 Supplement-1.** A.) Bar charts depicting the percentage of dead cells detected by propidium iodide (PI) staining in single cells treated with increasing concentrations of 2-DG for two hours. B.) Bar charts representing normalized HIV-1_JR-FL_ fusion ti vehicle in single cells as determined by the β-lactamase assay in cells treated with glucose-free medium for two hours before virus addition. C.) Gating strategy to detect single, live TZM-bl cells to quantify relative PerCP-Cy5 (CD4) and Alexa-Fluor 488 fluorescence to determine relative surface CD4 and CCR5 expression levels during listed treatment conditions (depicted by histograms shown to the right and bar charts shown in the bottom panels). D.) Representative fluorescence series of images (left) and single particle tracking traces (right) of Gag-eGFP (green) and DiD (red) dual-label HIV-1_JR-FL_ pseudovirions in TZM-bl cells (scale bar) illustrating that in control conditions (i.e. no 2-DG) that HIV-1_JR-FL_ entry in TZM-bl cells proceeds with a precipitous loss of Gag-eGFP and maintenance of DiD signal, indicative of endocytosis as previously described. E.) Representative fluorescence series of images (left) and single particle tracking traces (right) of Gag-eGFP (green) and DiD (red) dual-label HIV-1_JR-FL_ pseudovirions in TZM-bl cells (scale bar) illustrating that in 2-DG-treated conditions (i.e. no 2-DG) that HIV-1_JR-FL_ entry in TZM-bl cells may also proceed with a slow decay of DiD signal and maintenance of eGFP-Gag signal, indicative of hemifusion. *p<0.5, **p<0.01 *** p<0.001 as determined by either one-way ANOVA or one-way student’s T-test.

**Figure 4 Supplement-1.** A.) Bar charts depicting normalized Filipin MFI / cell in various treatment conditions listed to illustrate that 2-DG treated cells have plasma membrane cholesterol content similar to 1mM MBCD-treated cells and that supplementation with increasing concentrations of water-soluble cholesterol in 2-DG treated conditions rescues filipin staining signal. B.) Bar charts depicting filipin mean fluorescence intensity per MT4 T cell normalised to vehicle, indicating two hour incubation of increasing concentrations of oligomycin do not alter surface plasma membrane cholesterol. *p<0.5, **p<0.01 *** p<0.001 as determined by either one-way ANOVA or one-way student’s T-test.

**Figure 5 Supplement-1.** A.) (Left) Cartoon diagram depicting a model of the planarizable, push-pull membrane order probe Flipper-TR, constructed from two large dithienothiophene flippers, in liquid-disoreded (Ld) and liquid-ordered (Lo) environments. (Right) Average long lifetime (τ_1_) per TZM-bl cells under listed treatment conditions (n=30/condition). B.) Bar charts depicting the four components extracted from double-fitted photon lifetime histograms: τ_1_, τ_2_, τ_intensity_, and τ_amplitude_ of FliptR as a function of cell treatment. C.) Representative images of FlipTR stained cells in untreated conditions depicting the homogenous distribution of lifetimes throughout the entirety of the cell. D.) Raw-fits of TCSPC histograms (left) and normalized-fit to percentage of maximum total photon counts (right) of TCSPC FLIM images obtained from FlipTR-stained cells in specified treatment conditions. In addition, for reference, the associated long lifetimes (i.e. τ_1_) extracted from each image’s fit are listed. E.) (Left) Cartoon diagram model depicting membrane tension FRET probe MSS and its control KMSS probe (insensitive to membrane tension) KMSS. (Right) Representative pseudocoloured FRET efficiency (i.e. Ypet/eCFP) images of single TZM-bl cells expressing MSS under listed treatment conditions. F.) (Left) Dot plots depicting donor CFP lifetimes of the tension-sensitive MSS probes extracted from single cells in the treatment conditions listed. (Right) Dot plots depicting donor CFP lifetimes of the tension-insensitive KMSS probes extracted from single cells in the treatment conditions listed. *p<0.5, **p<0.01 *** p<0.001 as determined by either one-way ANOVA or one-way student’s T-test.

**Figure 5 Supplement-2.** A.) Compiled MSS lifetimes extracted from whole cells compiled from acquisitions of multiple viral fusion events (green) compared to 2-DG treated cells where viruses were incapable of fusion (blue) or rescued events due to cholesterol treatment (purple). B.) Bar charts comparing the MSS lifetimes of localised regions overlapping with mCherry-labelled HIV-1_JR-FL_ during virion fusion (i.e. mCherry signal loss) with virus-free control areas in the same cell at the timepoint of fusion. C.) Bar charts comparing the MSS lifetimes of localised regions overlapping with mCherry-labelled HIV-1_JR-FL_ (i.e. mCherry signal) with virus-free control areas in the same cell at the timepoint before fusion. D.) Bar charts comparing the MSS lifetimes of localised regions overlapping with regions where mCherry-labelled HIV-1_JR-FL_ fused with the host cell (i.e. mCherry signal lost) with virus-free control areas in the same cell at the timepoint after fusion. *p<0.5, **p<0.01 *** p<0.001 as determined by one-way student’s T-test.

