## Supplementary figures and images for "Single-cell glycolytic activity regulates membrane tension and HIV-1 fusion"

### Supp Files

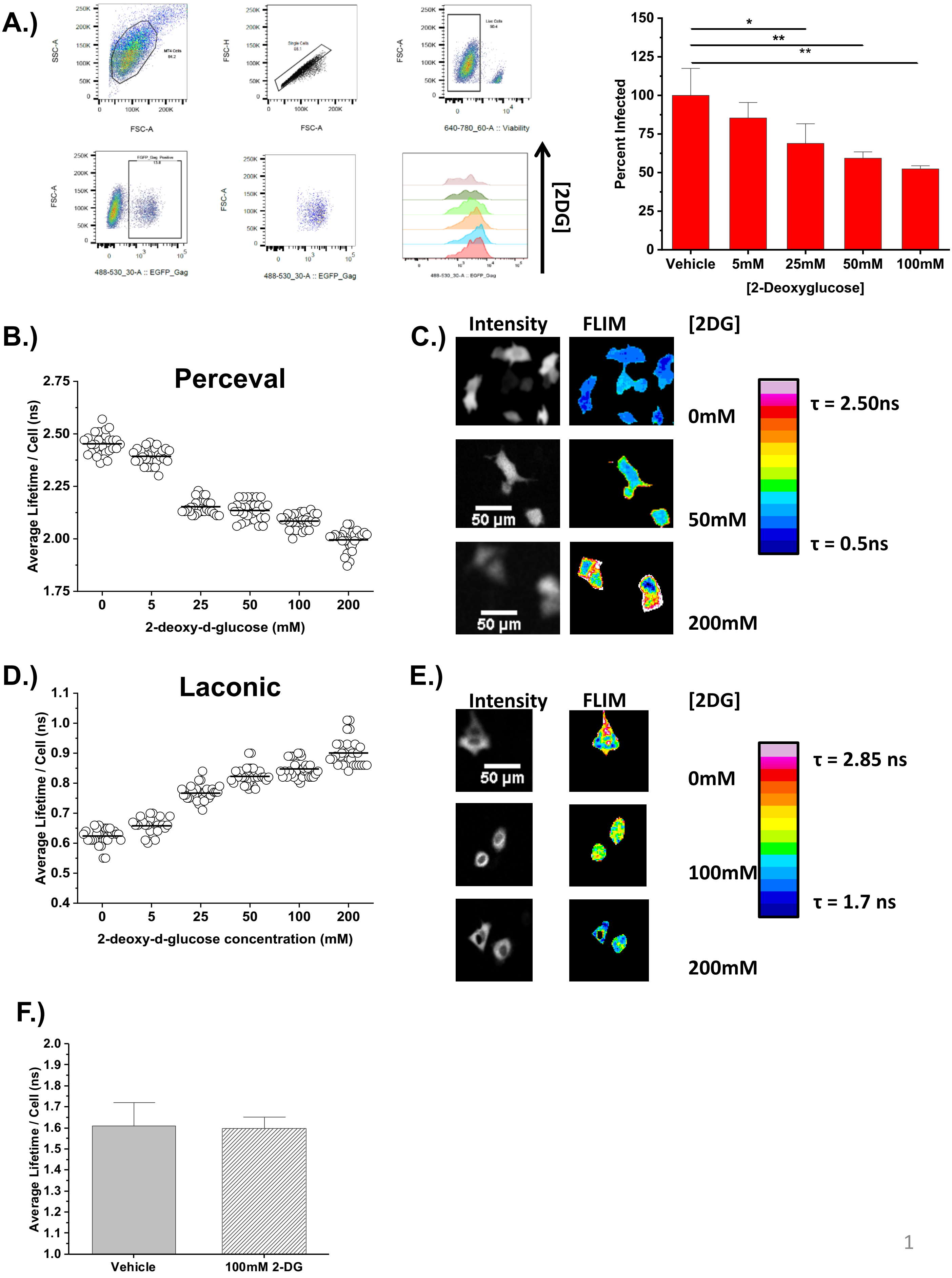

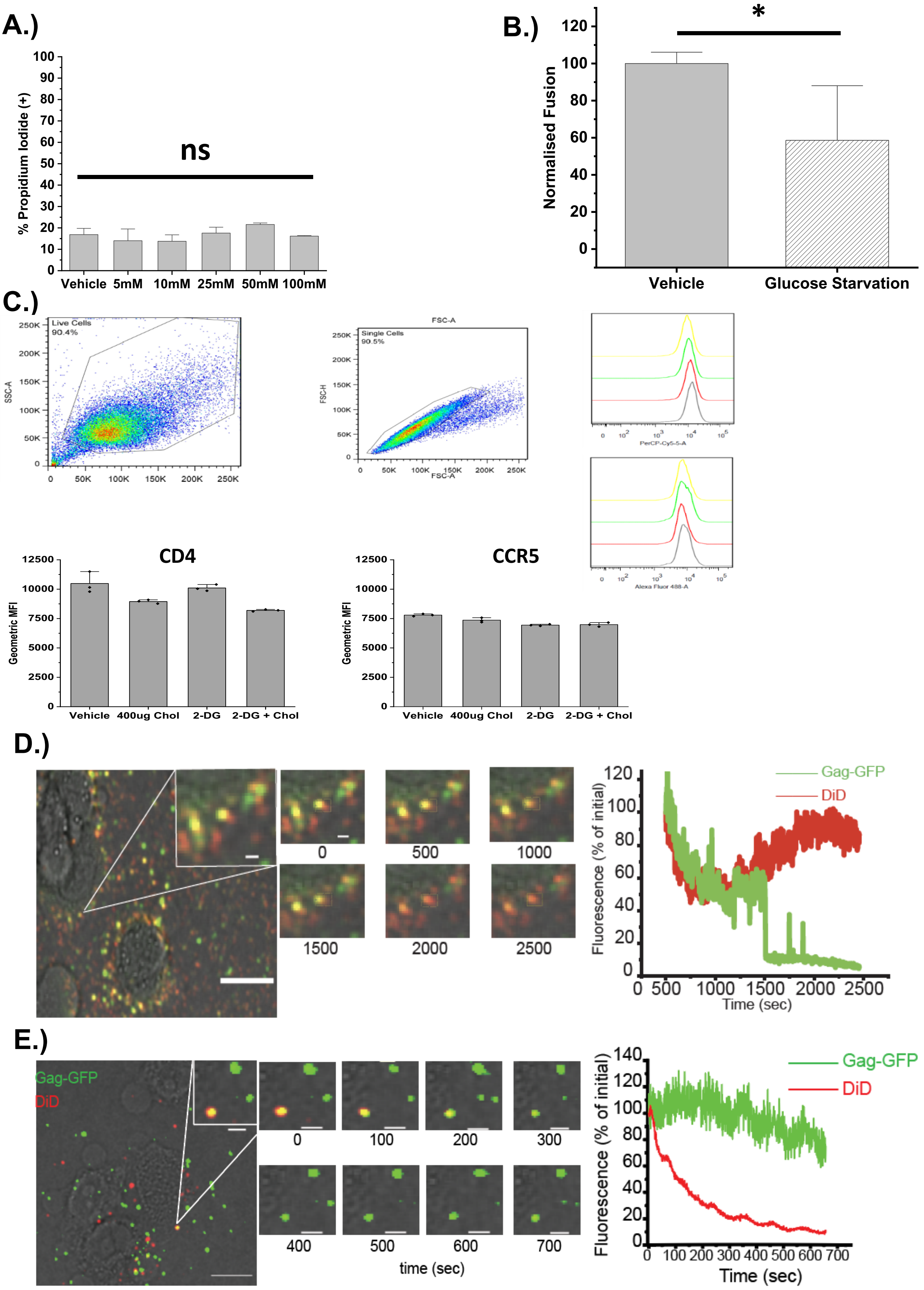

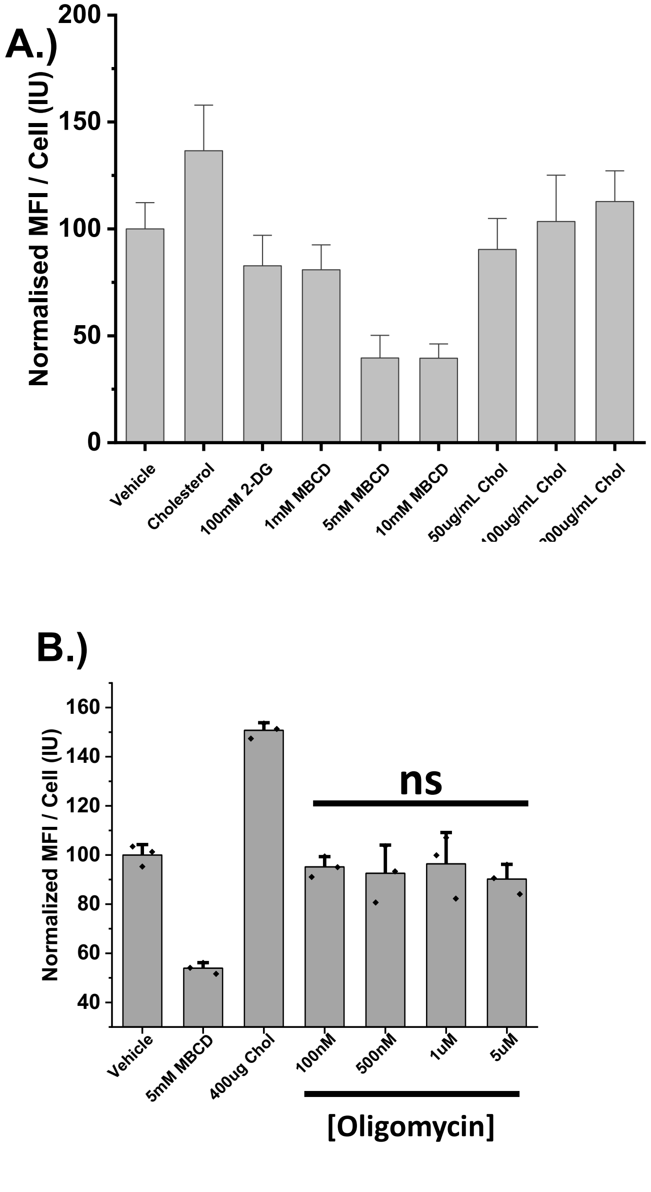

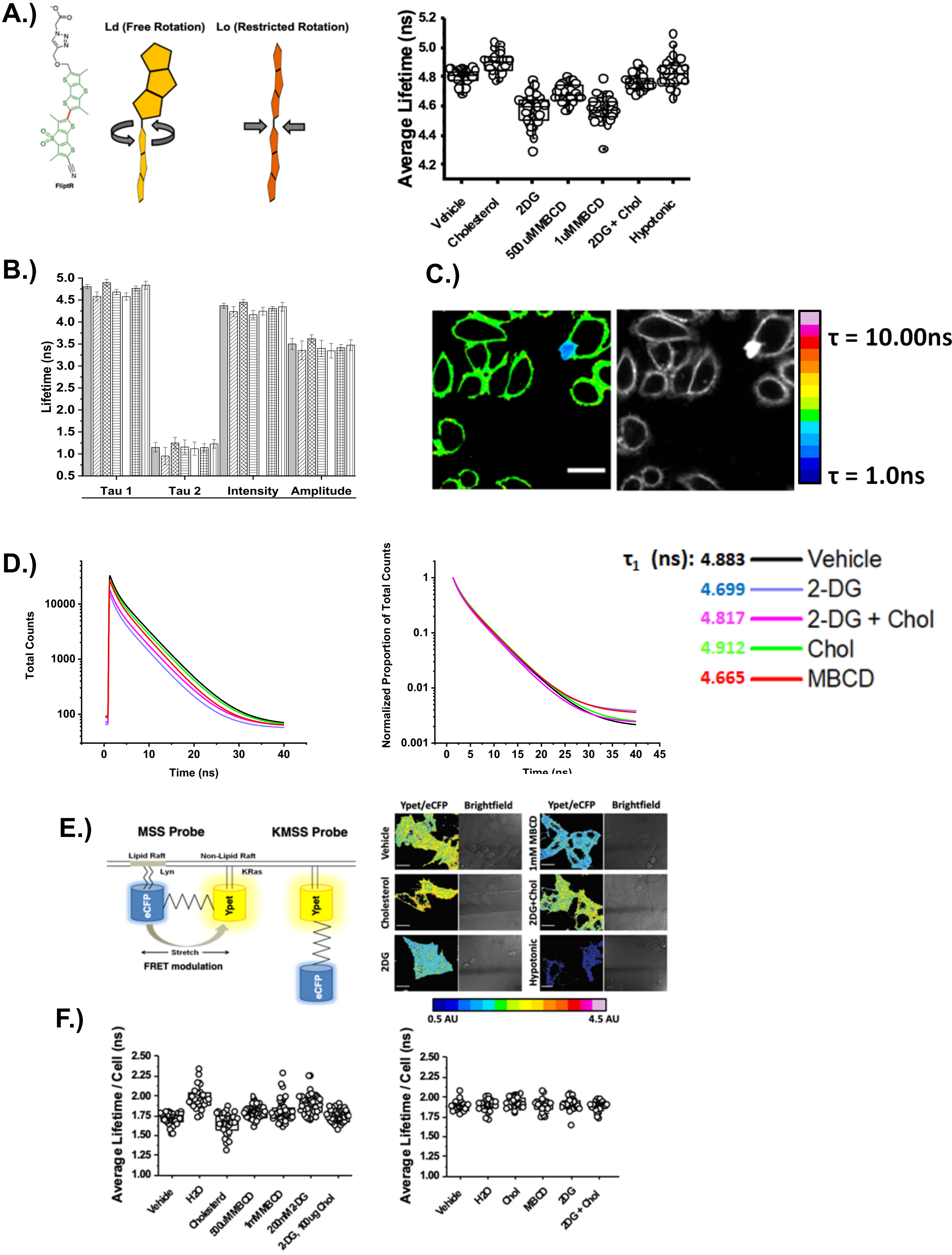

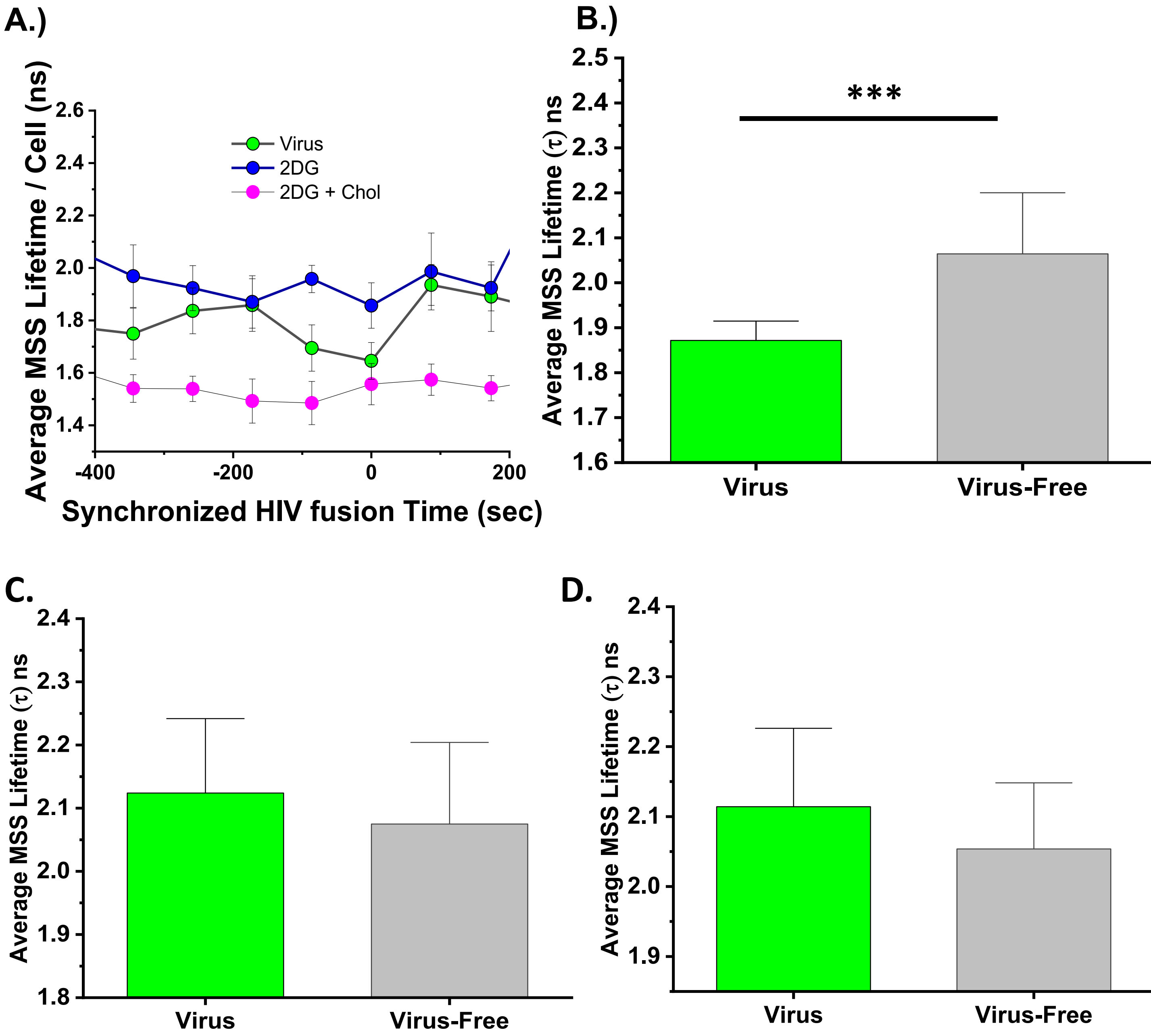
